# Combining next-generation sequencing and mtDNA data to uncover cryptic lineages of Mexican highland frogs

**DOI:** 10.1101/153601

**Authors:** Eugenia Zarza, Elizabeth M. Connors, James M. Maley, Whitney L.E. Tsai, Peter Heimes, Moises Kaplan, John E. McCormack

## Abstract

Recently, molecular studies have uncovered significant cryptic diversity in the Mexican Highlands, leading to the description of many new endemic species. DNA approaches to this kind of species discovery have included both mitochondrial DNA (mtDNA) sequencing and multilocus genomic methods. While these marker types have often been pitted against one another, there are benefits to deploying them together, as linked mtDNA data can provide the bridge between uncovering lineages through rigorous multilocus genomic analysis and identifying lineages through comparison to existing mtDNA databases. Here, we apply one class of multilocus genomic marker, ultraconserved elements (UCEs), and linked mtDNA data to a species complex of frogs (*Sarcohyla bistincta*) found in the Mexican Highlands. We generated data from 1,891 UCEs, which contained 1,742 informative SNPs for *S. bistincta* and closely related species and captured mitochondrial genomes for most samples. Genetic analyses based on both whole loci and SNPs agree there are numerous distinct and divergent lineages within *S. bistincta*. The SNP-based species tree provides the most conservative estimate of 8 well-supported lineages in three major clades. Having linked mtDNA data allowed us to tap into the large number of mtDNA sequences available on GenBank and identify one of these lineages as an already-described species, *S. pentheter*. One identified clade (containing 2 of the 8 lineages) was 10% divergent in mtDNA and paraphyletic with other *S. bistincta*, making this clade a clear candidate for species status. Phylogenies from UCEs and mtDNA mostly agreed in their topologies, but differed in that mtDNA suggested a more complex evolutionary history perhaps influenced by gene flow between some neighboring lineages. Our study demonstrates that the Mexican Highlands still hold substantial undescribed diversity. Combining multilocus genomic data with linked mtDNA data is a useful approach for identifying potential new species and associating them with already described taxa, which is especially important in groups with undescribed subadult phenotypes, where geographic ranges are unclear, or where phenotypes are conserved.

## Introduction

The Mexican Highlands are a global biodiversity hotspot (Myers et al. 2000). Recent molecular studies have uncovered significant cryptic diversity in the Mexican Highlands, leading to the description of new endemic species or the elevation of former subspecies to species rank (Bryson et al. 2010; Bryson et al. 2012; Bryson et al. 2011; McCormack et al. 2008; Rovito et al. 2015). At the same time, habitat loss threatens many of these species before they have even been described (Ponce-Reyes et al. 2012). Amphibians are particularly sensitive to habitat alterations, and many are threatened by habitat loss and invasive diseases (Stuart et al. 2008). Because most amphibians have reduced dispersal, they also show patterns of microendemism (Parra-Olea et al. 2014), meaning that important pockets of diversity and new species are still being uncovered in the Mexican Highlands (Campbell et al. 2009; Meik et al. 2006).

We investigate potential cryptic diversity within the *Sarcohyla bistincta* species complex in the Mexican Highlands. The genus *Sarcohyla*, which is considered distinct from *Plectrohyla* by some authors to reflect those species west of the Isthmus of Tehuantepec (Duellman et al. 2016), contains 24 species of stream-dwelling frogs, many of them critically endangered and many that have never been seen after their original discovery (references compiled in Stuart et al. 2008). Some species are thought to be in serious decline or extinct (Lips et al. 2004). The actual number of species, their relationships, and geographic ranges are not well known (Duellman 2001; Duellman et al. 2016; Faivovich et al. 2009). *S. bistincta* is the most broadly distributed member of the genus. It occurs in several mountain ranges separated by lowland barriers and might therefore be comprised of multiple species.

We sought to assess potential cryptic diversity in *S. bistincta* using multilocus genomic markers collected via next-generation sequencing (NGS) as well as mitochondrial DNA (mtDNA) data. Often, mtDNA and multilocus nuclear markers have been pitted against one another in biodiversity discovery (Edwards & Bensch 2009; Moritz & Cicero 2004; Zink & Barrowclough 2008). Mitochondrial DNA offers many benefits, including economy, efficiency, and wide comparative potential across species and studies. Consequently, there is now a vast trove of mtDNA sequences on GenBank. A drawback of using only mtDNA is that a single marker will often fail to accurately depict the speciation process (Edwards & Bensch 2009). In response, multilocus methods have multiplied (Edwards 2009; Fujita et al. 2012). While multiple loci help model a more realistic speciation process, multilocus studies suffer from the lack of a standardized marker set, which limits the ability to link uncovered lineages with species already identified and described in prior studies and in public databases. This is especially important in groups with multiple subadult phenotypes (e.g., insects and frogs) and where adult phenotypes are conserved across species. While many studies have used both types of markers, few have explicitly explored the benefits of linked mtDNA and NGS data at the level of the individual for lineage discovery and identification.

For our multilocus genomic markers, we use ultraconserved elements (UCEs). UCEs are an appealing multilocus marker set because the same loci are found across major branches of the tree of life, where they act as anchors for variable DNA in flanking regions (Faircloth et al. 2012). For instance, it is possible to capture the same set of 1000 or more UCEs across all mammals (McCormack et al. 2012), all reptiles (Crawford et al. 2012), or hundreds of UCEs across arachnid lineages separated by hundreds of millions of years (Starrett et al. 2016). While the power of UCEs for deep-level systematics is clear, their utility at shallower scales around the species level is still coming into focus (McCormack et al. 2016; Smith et al. 2014; Zarza et al. 2016). An added benefit of the UCE enrichment process (and all so-called “sequence capture” methods) is that whole mtDNA genomes are often captured as off-target “bycatch” (do Amaral et al. 2015), meaning each individual has associated nuclear and mtDNA data (e.g., Zarza et al. 2016). Our specific goals were to determine if UCEs are useful for addressing potential cryptic diversity within a species complex of frogs. Then, we assess whether linked mtDNA data, by providing a bridge to Genbank data, allows for more refined conclusions about the identification of any discovered lineages.

## Methods

### Sampling and Ingroup Determination

MK and PH collected tadpoles from January to June 2004 across most of the range where *Sarcohyla bistincta* are known to exist (Duellman 2001) in the Transvolcanic Belt of Michoacán, Morelos, and the state of México, the Sierra Madre del Sur of Guerrero, and the highlands of Oaxaca stretching into Puebla and Veracruz (Fig. 1; Table 1). Unsampled parts of the *S. bistincta* range include the far west Transvolcanic Belt in Michoacán and Jalisco, the far northwest in the Sierra Madre Occidental (Nayarit, Durango, and Sinaloa), and the far northeast in Hidalgo (see Fig. S1 for sampled and unsampled locations and known ranges of all *Sarcohyla* species).

**Table 1.**
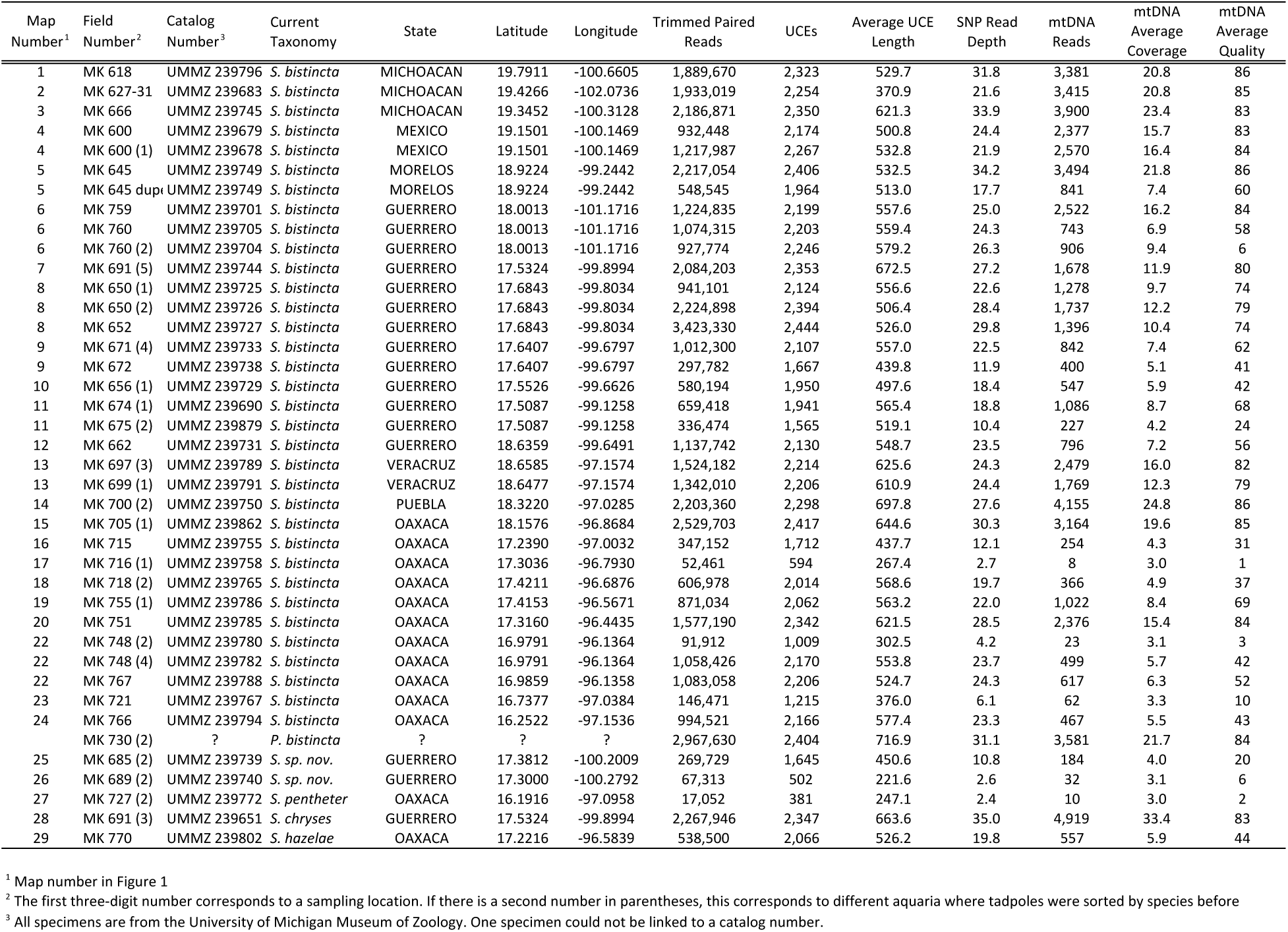
Specimen information and summary statistics for *Sarcohyla bistincta* and closely related species.

**Figure 1.**
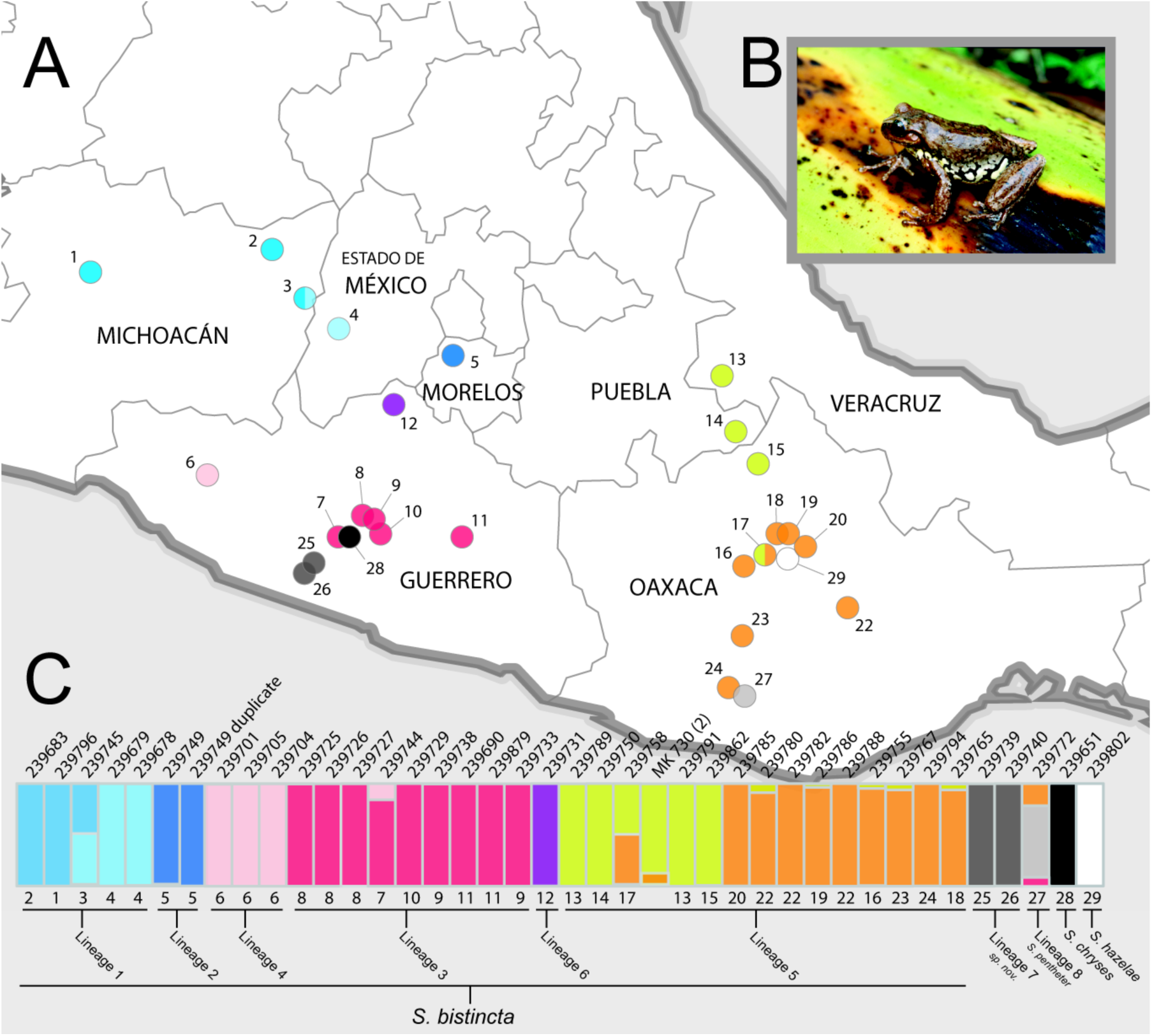
(A) Map of central Mexico showing sampling sites for *S. bistincta* and close outgroups, with numbers matching localities listed in Table 1 and colors matching Structure results below. Unsampled parts of the distribution of *S. bistincta* are shown in Fig. S1; (B) *S. bistincta* individual from near site 1; (C) Composite results of repeated Structure runs at *K* = 2 showing the finest detectable structure in the genetic data. Each vertical line represents an individual labeled with its UMMZ catalog number above and, in descending order below, the site number and the lineage number based on the SNAPP analysis. One tissue voucher MK 730 (2) could not be linked to a specimen voucher and thus its geographic locality is unknown.

Tadpoles were targeted to improve sampling efficiency, which allowed for a larger sampling range and sample density. After collection from a sampling location with a dip net, tadpoles were, to the extent possible, separated by species based on morphology and reared to subadults in the laboratory prior to vouchering. Species identification was based on the most recent diagnosis of *S. bistincta* and other closely related species (Duellman 2001). One tadpole was chosen for the tissue voucher, while the other individuals became physical vouchers with museum catalog numbers. Thus, we provide both field numbers and catalog numbers in Table 1 to provide a link to both the exact genetic material (field number) and the associated phenotype voucher representing that genotype (museum catalog number). Before limiting our taxonomic sampling to 38 individuals in a clade thought to represent *S. bistincta* (as we discuss later, some turned out to be already-described species nested within *S. bistincta*), we ran preliminary phylogenetic analyses including broader sampling of 45 *Sarcohyla* individuals and an outgroup genus *Exerodonta* to ensure we had correctly identified the ingroup (Table S1; Fig. S2).

### Sequence capture and next-generation sequencing

We extracted genomic DNA from tissue using a Qiagen (Valencia, CA) DNAeasy Blood and Tissue extraction kit. We visualized extractions on an agarose gel to ensure fragments were larger than 200 base pairs (bp) and quantified the resulting double-stranded DNA using a Qubit 2.0 Fluorometer (Carlsbad, CA). For each sample, we sheared 100 µl of 20 ng/µl concentration DNA to a size distribution with its peak between 400 and 600 bp using a Bioruptor ultrasonicator (Diagenode). We prepared libraries for each sheared sample with a KAPA (Boston, MA) LTP library preparation kit for the Illumina platform, attaching custom indexing tags (Faircloth & Glenn 2012) to each sample to allow sample pooling.

We enriched pools of eight samples using a set of synthetic RNA probes that target 5,060 tetrapod UCEs (MYbaits_Tetrapods-UCE-5K kit, Mycroarray) following the standard UCE enrichment protocol (Faircloth et al. 2012) with one modification. Amphibians have large and variable genome sizes with a high percentage of repetitive DNA (Olmo 1991). While we do not have information about the genome size and composition of *Sarcohyla* specifically, we wanted to decrease the potential risk of the probes hybridizing to repetitive elements (McCartney-Melstad et al. 2016). We thus increased by 6X the amount of the Cot-1 blocker, a synthetic DNA derived from chicken that binds to repetitive regions. After enrichment and recovery PCR, we verified the library size range with an Agilent 2100 Bioanalyzer (Palo Alto, CA). We quantified the enriched pools using qPCR and combined them in equimolar ratios before sequencing on an Illumina HiSeq 2000 lane (100-bp paired-end cycle) at the University of California Santa Cruz Genome Technology Center.

### Bioinformatics of next-generation sequencing data

We demultiplexed the Illumina raw reads and converted them to FASTQ format with the program bcl2fastq v.1.8.4 (Illumina, Inc.). To eliminate adapter contamination and low quality bases, we trimmed the FASTQC output with illumiprocessor (Faircloth 2013). We trimmed and assembled these reads into contigs with Trinity (Haas et al. 2013) and ABySS (Simpson et al. 2009), both of which are built into the PHYLUCE pipeline (Faircloth 2015). PHYLUCE uses LASTZ (Harris 2007) to align all assembled contigs to UCE probes in order to isolate only UCE contigs and to identify and eliminate paralogs.

### Phylogenetic trees from concatenated UCE data

We extracted UCE contigs into a single FASTA file and aligned the output for each locus using MAFFT (Katoh et al. 2005). We required that 75% of the taxa needed to have data for a given locus to be included in the final concatenated matrix, which led to dropping loci that did not meet this threshold. We then constructed a maximum-likelihood (ML) tree in RAxML v8.0.19 (Stamatakis 2014) under the GTRGAMMA model of evolution with 100 bootstrap searches, followed by a search for the tree with the highest likelihood.

### Mitochondrial DNA assembly and analysis

We identified and assembled mtDNA genomes from off-target, trimmed Illumina reads using the reference genome of a closely related species, *Hyla annectans* (Genebank accession number KM271781; Ye et al. 2016). We used MITObim 1.7 (Hahn et al. 2013), a Perl wrapper for MIRA 4.0.2 (Chevreux et al. 1999), that takes an iterative mapping approach for assembly. We conducted *de novo* annotation of the assembled mtDNA regions with the MITOchondrial genome annotation Server, MITOS (Bernt et al. 2013). We selected for analysis only those individual genomes with MIRA quality score greater than 30. We aligned each protein-coding region separately in Geneious vR8 (Kearse et al. 2012) using MUSCLE (Edgar 2004) and corrected the alignments manually when necessary and constructed a concatenated mtDNA matrix, which we also ran in RAxML v8.0.19.

We melded this mtDNA data with existing *Sarcohyla* and *Plectrohyla* mtDNA data on Genbank to determine whether any of the lineages we uncovered in *S. bistincta* relate to already-described species. We determined that *cytochrome b* is the best-represented mtDNA gene on Genbank for this group. We downloaded all existing *cytochrome b* sequences from *Sarcohyla* and *Plectrohyla* taxa. We combined these sequences with those from a subset of our *S. bistincta* samples, choosing the individual with the most raw reads from each major genetic lineage in the UCE tree. We aligned the trimmed, filtered reads for each individual to a *Sarcohyla cytochrome b* reference sequence and formed a consensus sequence for each individual from the mapped reads. We then created an alignment and generated a phylogeny using BEAST v2.4.2 (Bouckaert et al. 2014). We repeated this process with 16S data because we suspected *S. pentheter* was closely related to our ingroup, but *S. pentheter* had no *cyt b* sequence on Genbank.

### Calling SNPs from UCE loci

We called SNPs from UCE loci so that we could run genetic clustering tests and infer a species tree using SNAPP (Bryant et al. 2012), which uses SNPs as input data. Calling SNPs requires a reference sequence, and we chose the sample with the most UCE contigs recovered within the ingroup (UMMZ 239727). We then used BWA (Li & Durbin 2009) to map the reads of each sample to this reference. We used SAMtools (Li et al. 2009) to sort the reads, and Picard (http://broadinstitute.github.io/picard) to identify and remove PCR duplicates. We realigned the mapped reads to minimize mismatched bases due to indels, and we removed indels using the Genome Analysis Toolkit 3.2 (GATK; McKenna et al. 2010) and a custom script (indelrealigner.sh), as suggested by the Best Practices workflow (DePristo et al. 2011; van der Auwera et al. 2013).

There is no SNP database available for treefrogs, so we followed best practices for base recalibration for non-model organisms suggested by GATK (McKenna et al. 2010). This consists of (1) doing an initial round of calling SNPs on the original, uncalibrated data, (2) selecting the SNPs with the highest confidence (a minimum emission and call quality of 40 or more), and (3) using these SNPs as the reference database of known SNPs. We executed four rounds of base recalibration on the original data to filter out systematic error using a custom script. We called genotypes on the last recalibrated BAM file. We used vcf-tools (Danecek et al. 2011) to select one SNP per UCE and produce two data sets, one allowing 25% missing data for STRUCTURE v 2.3.4 (Pritchard et al. 2000), and one with no missing data – a requirement for SNAPP as implemented in BEAST v2.2.1 (Bouckaert et al. 2014).

### STRUCTURE analyses

We used STRUCTURE v2.3.4 (Pritchard et al. 2000) as an unbiased way to assess the limits of fine-scale genetic structure in our data. Our intent with using STRUCTURE was not to determine the single most likely number of genetic clusters; thus, we did not use a method for identifying the “true K” (Evanno et al. 2005), which can underestimate fine-scale population structure (Janes et al. 2017). Rather, our goal was to determine the maximum possible number of genetic clusters in our data (e.g., Brown et al. 2007; Gowen et al. 2014). We began by analyzing all individuals of *S. bistincta* plus two outgroup species *S. chryses* and *S. hazelae* under *K* = 4, reasoning that this would likely split out the two outgroups as well as reveal at most one division within *S. bistincta*. After this, each identified genetic cluster within *S. bistincta* was further analyzed at *K* = 2 until no coherent geographically-based structure was evident in the plots. All runs were completed twice and each used an admixture model and 10M generations with 1M generations as burn-in, which led to convergence for all analyses.

### SNAPP tree and species delimitation

We generated a species tree from the SNP matrix using SNAPP 1.1.10 (Bryant et al. 2012). This analysis included putative *S. bistincta* samples (again, later shown to include other species nested within) and one outgroup, *S. chryses*. For this run, we made no *a priori* assumptions about how individuals grouped into species and allowed each individual to be considered its own “species” (i.e., terminal tip). We ran two instances of SNAPP for seven million generations using default priors. We combined tree and parameter files from both runs with LogCombiner 2.1.3 and displayed the full set of likely species trees with Densitree v2.2.1 (Bouckaert et al. 2014).

## Results

### NGS summary statistics

Detailed summary statistics for each of the 38 ingroup samples and two outgroups are described in Table 1. ABySS produced longer contigs than Trinity, and a higher number of UCE loci, so we used ABYSS contigs in all downstream analyses. Reads per sample ranged from 17,052 to 3,423,330 with an average of 1,185,165 reads. The number of identified UCEs ranged from 381 to 2,444 with an average of 1,976 UCEs. The mean length of individual UCE loci per individual ranged from 222 to 717 bp with an average of 522 bp. On average, 18% of the assembled contigs corresponded to unique UCE loci.

For SNP calling, across 40 samples of *S. bistincta* and outgroups, 9% of the trimmed reads mapped to our designated reference individual. SNP read depth ranged from 2.4 to 35.0 with an average depth of 21.2. The recalibration and quality control steps resulted in an initial matrix of 16,578 SNPs. After removing non-biallelic loci, selecting one SNP for every UCE, and allowing 25% missing data, there were 1,742 SNPs left in the STRUCTURE data set, while the 100% complete data matrix for SNAPP contained 399 SNPs.

### UCE phylogeny from concatenated data

Our more taxonomically inclusive data set with all available *Sarcohyla* and outgroup *Exerodonta xera* (Table S1) contained 1,866 UCE loci and 1,030,450 bp for a concatenated analysis. The resulting ML tree (Fig. S2) showed strong support for monophyly of *Sarcohyla*, and identified *S. arborescandens* and *S. cyclada* as sister species that together form a clade sister to the rest of the *Sarcohyla* included in the study. We limited further analyses to a smaller data set of 40 samples with *S. hazelae* as the outgroup (Table 1). This focal data set contained 1,891 UCE loci and 1,038,600 bp.

The ML tree of these 40 samples found strong support for many clades within the species currently described as *S. bistincta*, conforming to distinct geographic areas (Fig. 2a).

**Figure 2.**
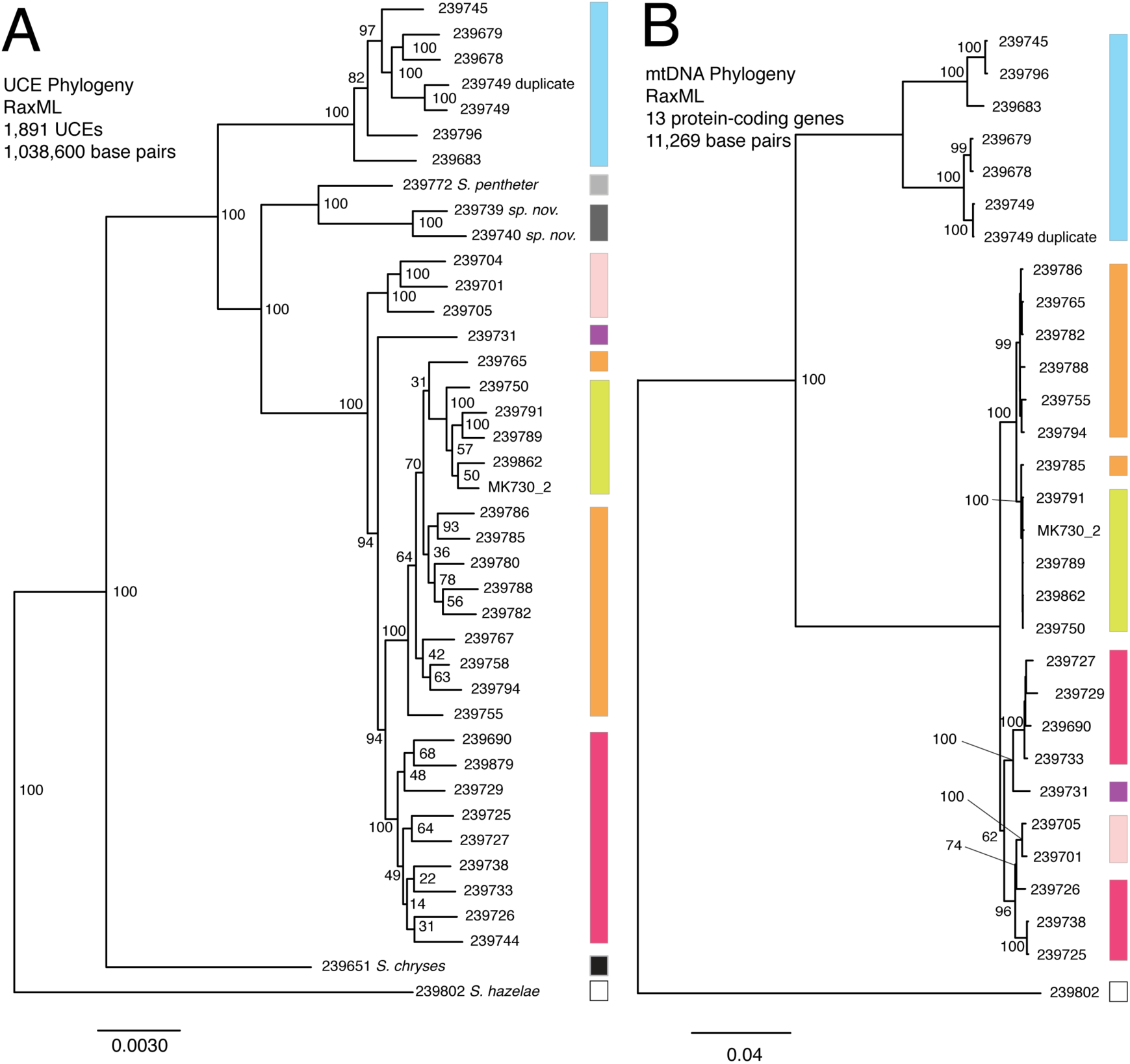
(A) UCE tree; (B) mtDNA tree. Colors match Structure groups identified in Fig. 1. Tips are labeled with their UMMZ catalog number.

In brief, there were three clades on relatively long branches: (1) a clade distributed across the Transvolcanic Belt (shaded blue in figures); (2) a clade inhabiting two disjunct areas along the coastal slopes of the Sierra Madre del Sur in Guerrero and Oaxaca (shaded gray); and (3) a clade broadly distributed in the Sierra Madre del Sur (shaded red and pink), the Oaxaca Highlands (shaded yellow and orange), and one individual in the southern portion of the Transvolcanic Belt (shaded purple). One individual that nested within *S. bistincta* was labeled as a different species, *S. mykter*, from Guerrero. We suspect that this sample was mislabeled and is actually a duplicate of an *S. bistincta* sample already included in the study because their field numbers are similar (last two digits transposed) and the two samples grouped together in all analyses. We have left this sample in all analyses, but have labeled it as a duplicate of *S. bistincta* UMMZ 239749. Another tissue voucher, MK 730 (2), could not be linked definitively to a physical voucher, and thus its geographic location is unknown. Its tip label has been left as the field number. Connecting UCE with mtDNA data, discussed later, revealed that one specimen in the gray clade (UMMZ 239772) is an already-described species, *S. pentheter*.

### mtDNA tree

Our final concatenated mtDNA matrix (individual *cyt b* and 16S tree are discussed later) was 11,269 base pairs including gaps, as coverage of the mtDNA genome varied from sample to sample in accordance with the non-targeted nature of the DNA collection (Table 1). Relationships in the ML tree (Fig. 2b) among the 29 individuals with high quality scores were similar to the concatenated UCE tree with two key differences, both within the broadly distributed clade in Guerrero and Oaxaca: (1) in the mtDNA tree, individuals from eastern and western Guerrero (shaded pink and red in figures) formed a clade, whereas they were more distantly related in the UCE tree; (2) in the mtDNA tree, individual UMMZ 239731 (shaded purple) was nested within the Guerrero clade instead of being sister to a much more inclusive clade, as in the UCE tree.

### Structure analysis

As expected, the first run of STRUCTURE at K = 4 split the two outgroup species into distinct clusters and split the remaining individuals into two clusters. Further analysis of each cluster at K = 2 revealed ten genetic clusters in all (Fig. 1c), which are largely concordant with clades observed in the UCE and mtDNA phylogeny. Most individuals did not share assignment among clusters, although a few individuals showed mixed assignment among clusters.

### SNAPP species tree

The SNAPP tree and its cloudogram of posterior species trees (Fig. 3) revealed eight well-supported lineages consistent with many of the genetic clusters in the Structure analysis and with relationships in the UCE and mtDNA whole-genome phylogenies. With respect to the discrepancies between UCE and mtDNA trees detailed above, the SNAPP tree agreed with the mtDNA tree that eastern and western Guerrero individuals form a clade, but agreed with the UCE tree in the placement of individual UMMZ 239731.

**Figure 3.**
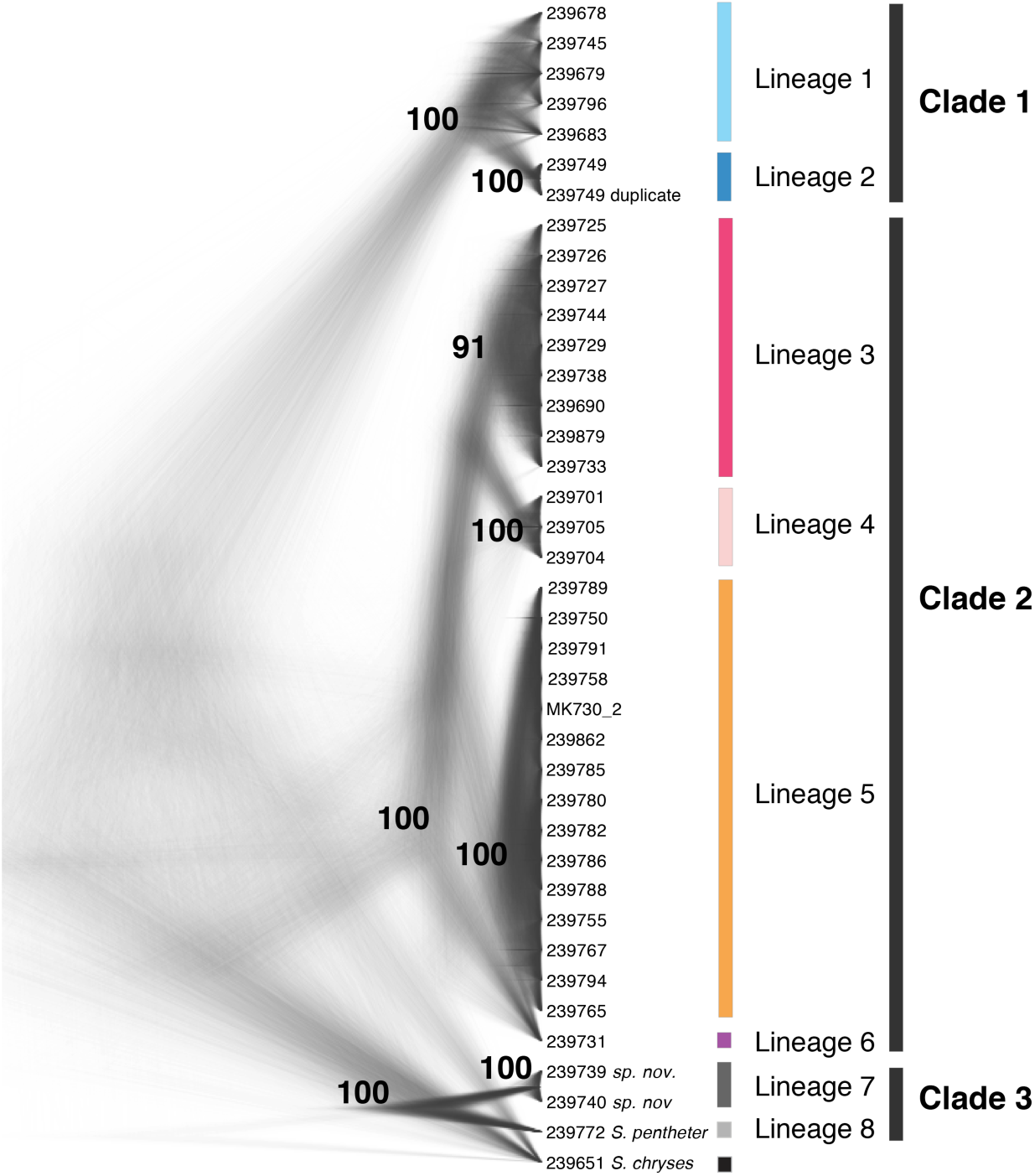
Cloudogram of the posterior distribution of SNAPP trees from 399 high-quality SNPs mined from UCE loci. Tip labels are UMMZ catalog numbers. Lineages and clades are discussed in text. Colors match genetic clusters from Figure 1.

### mtDNA phylogeny combining new data with Genbank sequences

Using 16S sequences, we determined that one lineage from Fig. 3 matched an *S. pentheter* sequence on Genbank. This individual (UMMZ 239772) had one of the lowest read counts of any samples and very few mtDNA reads. However, five Illumina reads mapped to 16S covering 421 bp of the 681 bp reference sequence (Genbank *S. pentheter* accession number DQ055825). Over this stretch, UMMZ 239772 was identical to the *S. pentheter* reference. As a point of comparison, UMMZ 239679 (a member of the blue *S. bistincta* Lineage 1 in the Transvolcanic Belt) had 70 differences across the 681 bp (10.3% divergence). This DNA identification of UMMZ 239772 as *S. pentheter* was later confirmed by re-examining the subadult specimen.

After confirming UMMZ 239679 as *S. pentheter*, we generated a Bayesian tree of *cytochrome b* combining the samples from this study with Genbank sequences (Fig. 4). This tree revealed not only that *S. pentheter* was nested within the current *S. bistincta*, but so was another species not included in our sampling, *S. calthula*. The tree also helped clarify relationships outside of *S. bistincta* by supporting *S. chryses* + *S. mykter* to be sister to the *S. bistincta* + *S. pentheter* + *S. calthula* clade.

**Figure 4.**
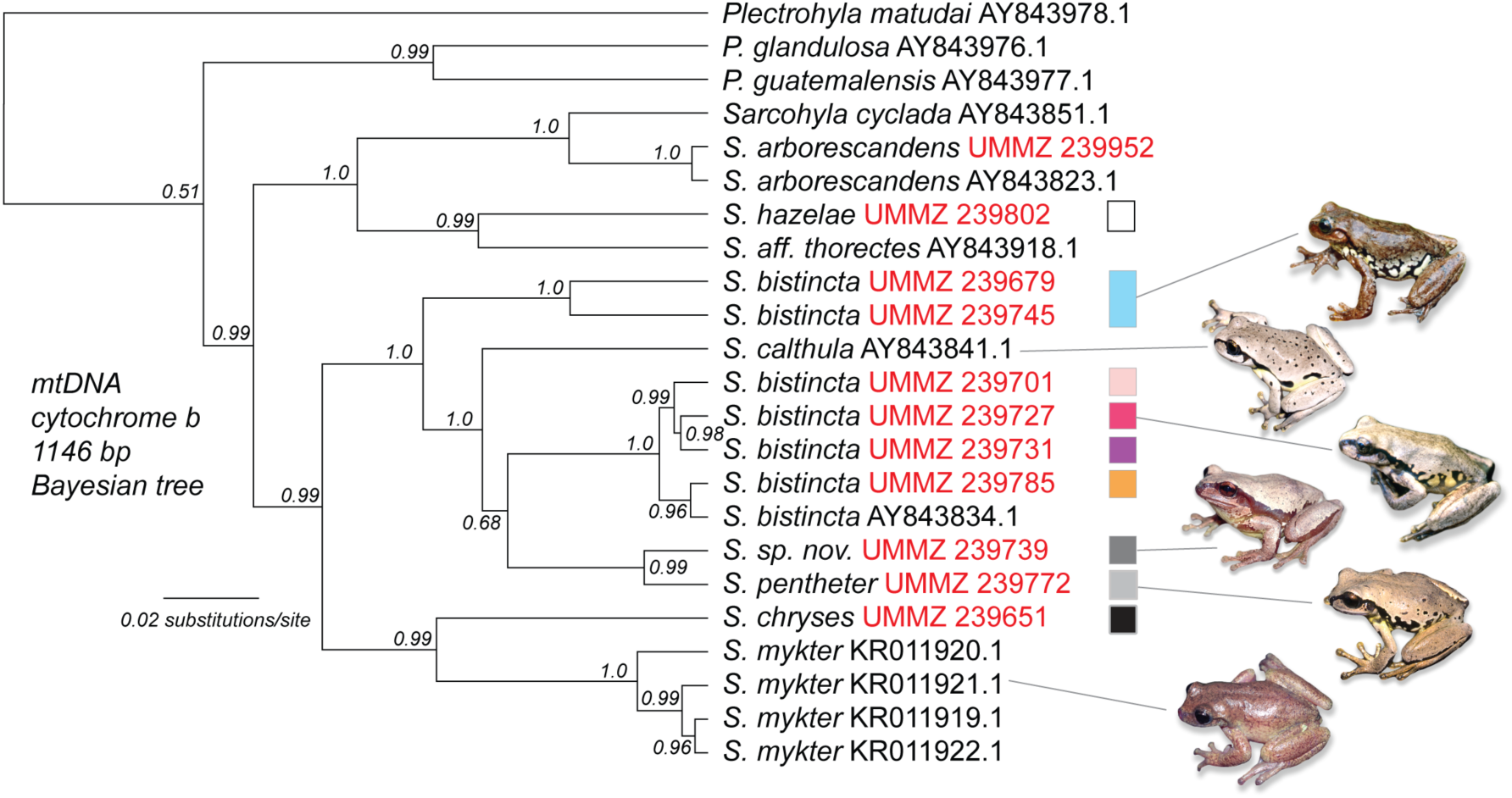
Phylogeny of mtDNA for a subset of the individuals from this study (labeled with red UMMZ catalog numbers) combined with existing sequences from Genbank (labeled with accession numbers). Colored boxes relate to genetic lineages in prior figures. The tree was rooted with *Exerodonta xera*. Photo credit: Peter Heimes.

## Discussion

### Bridging multilocus and mtDNA data for cryptic lineage discovery

Multilocus NGS data identified numerous divergent lineages within the *Sarcohyla bistincta* complex, supporting similar patterns observed in a recent study of this complex based on mtDNA and a few nuclear genes (Caviedes-Solis & Nieto-Montes 2017). We discuss these uncovered lineages in detail below, but note here that linking NGS with mtDNA data allowed us to query our lineages – first uncovered with multilocus genomic data – against GenBank to see if any of them corresponded to already-described species. Doing so revealed that *S. bistincta* is paraphyletic, with two already-described species nested within its current taxonomic limits (Fig. 4). The approach of discovering lineages with multilocus data and identifying them with the help of linked mtDNA data should be especially useful in understudied groups where basic natural history information is lacking.

### UCEs as a universal genomic marker set for species discovery?

Although it requires further study, UCEs could answer the call for an multilocus DNA marker set that satisfies criteria for both ease-of-use, universality, and genomic coverage (Coissac et al. 2016). UCE probe sets are now available for many taxonomic groups (Faircloth et al. 2015; Faircloth et al. 2013; Starrett et al. 2016). They capture a discrete and replicable portion of the genome, in this case a set of around 2,000 loci in frogs (from a larger set of ∼5,000 vertebrate loci) that query approximately 1,000,000 base pairs, or 0.02% of the frog genome. The replicable nature of UCEs sets them apart from other types of genomic markers, like RAD loci, which can vary from experiment to experiment (DaCosta & Sorenson 2014) and find fewer orthologs with increasing phylogenetic distance (Cruaud et al. 2014). Other alternatives to UCEs exist, like exons, but in mammals exons had fewer loci conserved over broad taxonomic scales than UCEs, making them less able to be universally applied (McCormack et al. 2012). A recent study successfully applied UCEs in conjunction with species delimitation methods to two frog genera, *Melanophryniscus* and *Brachycephalus* (Pie et al. 2017). As more studies are conducted, one future research avenue would be determine how much locus-sharing occurs among studies, and whether objective benchmarks for species-level divergence can be identified.

### Implications for species limits within Sarcohyla bistincta

The monophyletic group containing *Sarcohyla bistincta* is comprised of many distinct lineages as well as two already-described species (*S. pentheter* and *S. calthula*). These lineages are grouped into three fairly divergent clades, which agree with clades uncovered in a recent study based on fewer markers (Caviedes-Solis & Nieto-Montes 2017). In discussing these clades and lineages below, we use the most conservative estimate of 8 lineages, supported by the species tree, as a framework. Within that framework, we discuss the potential for further genetic structuring, as suggested by genetic clustering results and individual UCE and mtDNA phylogenies. We do not attempt to estimate the “true” number of species through species delimitation or other methods, because we feel species delimitation is best carried out through integrative taxonomy (Will et al. 2005), including at minimum analysis of the phenotype.

**Clade 1** – Transvolcanic Belt of central Mexico. This clade is sister to the rest of *S. bistincta* plus two other species, *S. pentheter* and *S. calthula*, and is 10% divergent in mtDNA from other *S. bistincta*, making it an obvious candidate for species status. This clade might itself contain multiple species in the form of the lineages below. Unsampled populations of *S. bistincta* in the Sierra Madre Occidental (Fig. S1) are very likely related to Lineage 1 of this clade, but should be included in future studies, as they might be distinctive.

<u>Lineage 1</u> (light blue in Fig. 1) – Michoacán to western Mexico state. The Structure results show fine-scale genetic structure across this range, and the presence of a geographic and genetic intermediate hints at continuity of gene flow along the distribution from sites 1 to 4 in Figure 1. In addition to unsampled populations mentioned above, some populations in far western Michoacán (Fig. S1) are also unsampled and could reveal further genetic structure.

<u>Lineage 2</u> (dark blue in Fig. 1) – Morelos. Denser sampling between sites 4 and 5 could help determine whether the genetic distinctness of this individual in the mtDNA and UCE trees is a true discontinuity or the result of a sampling gap.

**Clade 2** – Guerrero to Puebla and Veracruz and south through Oaxaca. This clade was also found to be distinct in a recent study (Caviedes-Solis & Nieto-Montes 2017) and forms the core *S. bistincta* (*sensu stricto*).

<u>Lineage 3</u> (red in Fig. 1) – Central and eastern Guerrero. Members of this lineage are distinct from Lineage 4 and are monophyletic in the UCE-based trees (though not in the mtDNA tree). Further sampling in between site 6 and site 7 would clarify whether the genetic discontinuity between Lineages 3 and 4 results from a sampling gap.

<u>Lineage 4</u> (pink in Fig. 1) – Western Guerrero. This lineage is monophyletic in all trees, although only a few individuals were sampled from a single locality.

<u>Lineage 5</u> (orange and yellow in Fig. 1) – Puebla, Veracruz, and Oaxaca. This lineage contains the type locality for *S. bistincta*. Central and southern Oaxaca individuals (orange) are genetically distinct from individuals to the north (yellow). One genetic intermediate in central Oaxaca suggests genetic continuity across this range. An unsampled northern population in Hidalgo is most likely related to this lineage, and should be included in future studies.

<u>Lineage 6</u> (purple in Fig. 1) – far northern Guerrero. This lineage is distinct but represented by only a single individual. However, in the mtDNA tree, this individual is nested within Lineages 4 and 5 above. Sampling more individuals is needed to determine how distinct this lineage might be.

**Clade 3** – Pacific slope of Guerrero and Oaxaca. This clade contains two species: one already described (*S. pentheter*) and one currently being described (*sp. nov*.; Kaplan et al. in prep). The relationship of *S. calthula* to this clade is unclear in our results, as bootstrap support was low in the combined mtDNA tree (Fig. 4). Caviedes-Solis et al. (2017) placed *S. calthula* sister to *S. pentheter* based on five genes. Thus *S. calthula* is likely also a member of Clade 3.

<u>Lineage 7</u> (dark gray in Fig. 1) – Pacific slope of Guerrero. This lineage was being described as a new species (we call it *sp. nov*.) on the basis of phenotypic differences before this genetic study was begun (Kaplan et al. in prep). Thus, our study lends support to species status for this lineage.

<u>Lineage 8</u> (light gray in Fig. 1) – *S. pentheter*. Pacific slope of Oaxaca.

The three clades and nearly all of the lineages were distinct in the mtDNA tree as well as the UCE tree (Fig. 2). The mtDNA tree, however, supports a more complicated history for Lineages 3, 4, and 6 in Guerrero. It is unclear why UCE and mtDNA results differed in this regard, but some reticulate processes might have influenced the mtDNA genomes of these lineages, perhaps, given their close geographic proximity, mtDNA capture of one lineage by another through gene flow (e.g., Bryson Jr et al. 2010).

Additional insights into broader *Sarcohyla* relationships offered by the mtDNA tree include support for a previously hypothesized close relationship between *S. hazelae* and *S. thorectes* (Caviedes-Solis & Nieto-Montes 2017; Faivovich et al. 2009), and a sister relationship between *S. mykter* and *S. chryses* (also found by Caviedes-Solis et al. (2017)). As *Sarcohyla* is very poorly represented by voucher specimens and DNA sequences (Fig. S1), a complete understanding of the history of this genus must await more complete taxonomic and genomic sampling. Unfortunately, there appear to be some microendemic *Sarcohyla* that might have already gone extinct (Lips et al. 2004), especially in the Oaxacan highlands, although recent resurveys provide hope for rediscoveries (Delia et al. 2013).

This and other recent studies show there is still substantial diversity remaining to be described in the Mexican Highlands. Genetic studies to uncover this diversity might use different approaches and marker types, but these efforts need not be in opposition. As our study shows, NGS and mtDNA data work well together, when lineages uncovered via multilocus methods (but with linked mtDNA data) can be checked against and combined with existing mtDNA databases like Genbank. Identification of vouchered material might not be a problem with some well-studied groups, like birds, but it is invaluable for groups with undescribed larval stages, unclear range boundaries, and difficult or highly conserved phenotypes. As the destruction of native habitats continues apace, it is important that we identify distinctive lineages and geographic centers of diversity before they are lost.

## List of abbreviations

UCEs: ultraconserved elements
SNPs: single nucleotide polymorphisms
mtDNA: mitochondrial DNA
bp: base pairs
ML: maximum-likelihood
UMMZ: University of Michigan Museum of Zoology
RAD loci: restriction digest-associated loci

## Declarations

### Ethics approval and consent to participate

This study complied with standard ethical guidelines for the rearing and collection of tadpoles.

### Consent for publication

Not applicable.

### Availability of data and materials

The datasets generated and analyzed in the current study will be made available on Genbank (BioProject ID PRJNA393258) and Dryad upon publication.

### Competing Interests

The authors declare that they have no competing interests.

### Funding

The project was partially funded by National Science Foundation grant DEB-1258205 to JM.

### Authors’ contributions

MK and PH collected samples in the field; MK and JEM designed the study; EZ and WLET produced the genomic data; EZ, EMC, and JMM analyzed the data; JEM and EZ wrote the manuscript; JEM created the figures; All authors read and approved the final manuscript.

## Acknowledgments

We thank Greg Schneider and Ron Nussbaum at UMMZ for providing access to tissue samples and vouchers, and Oscar Flores for help with permits.

## Tables

**Table S1.**
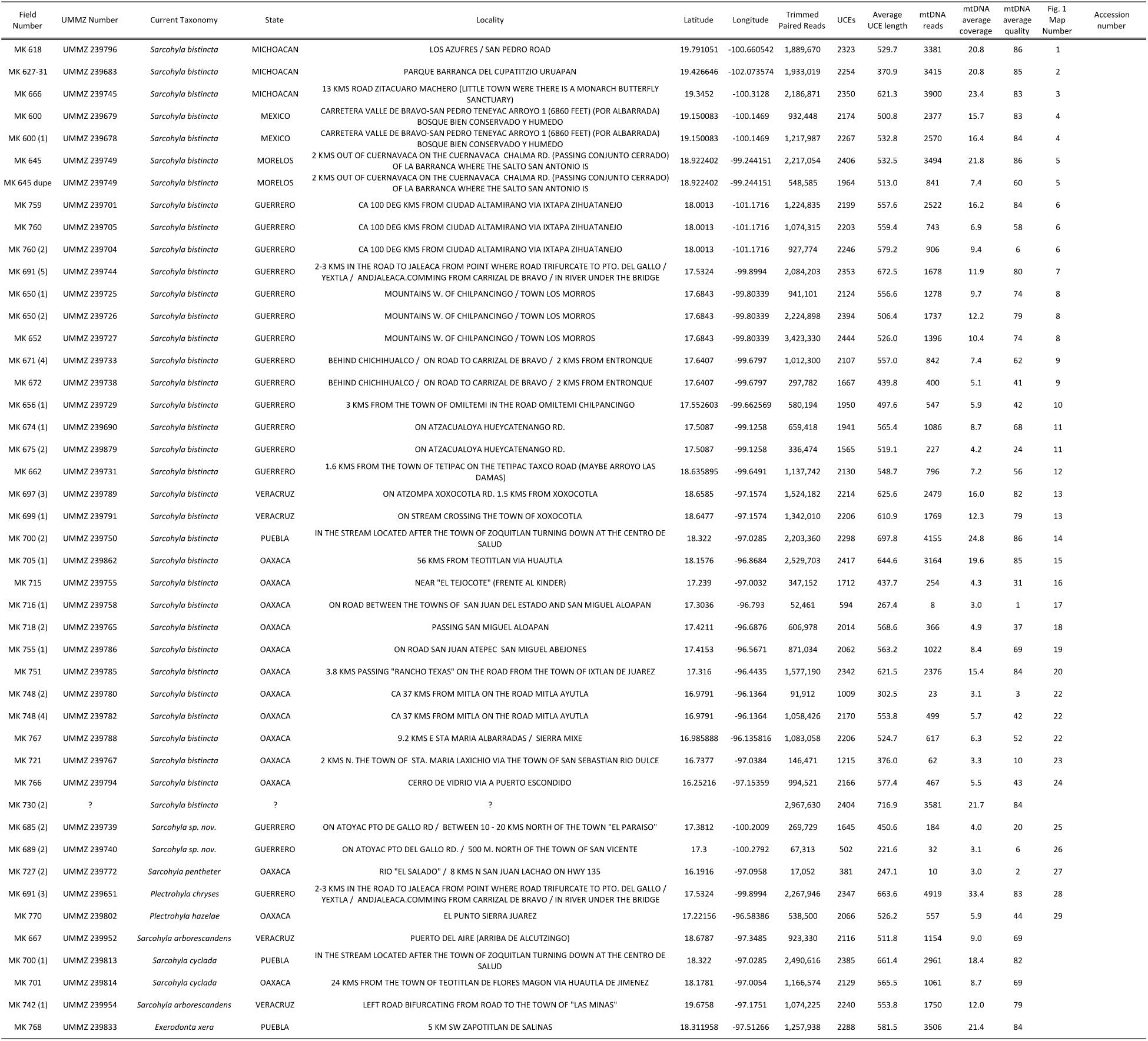
Information and summary statistics on all 45 samples used to determine the ingroup for this study.

## Additional Files

**Fig. S1.**
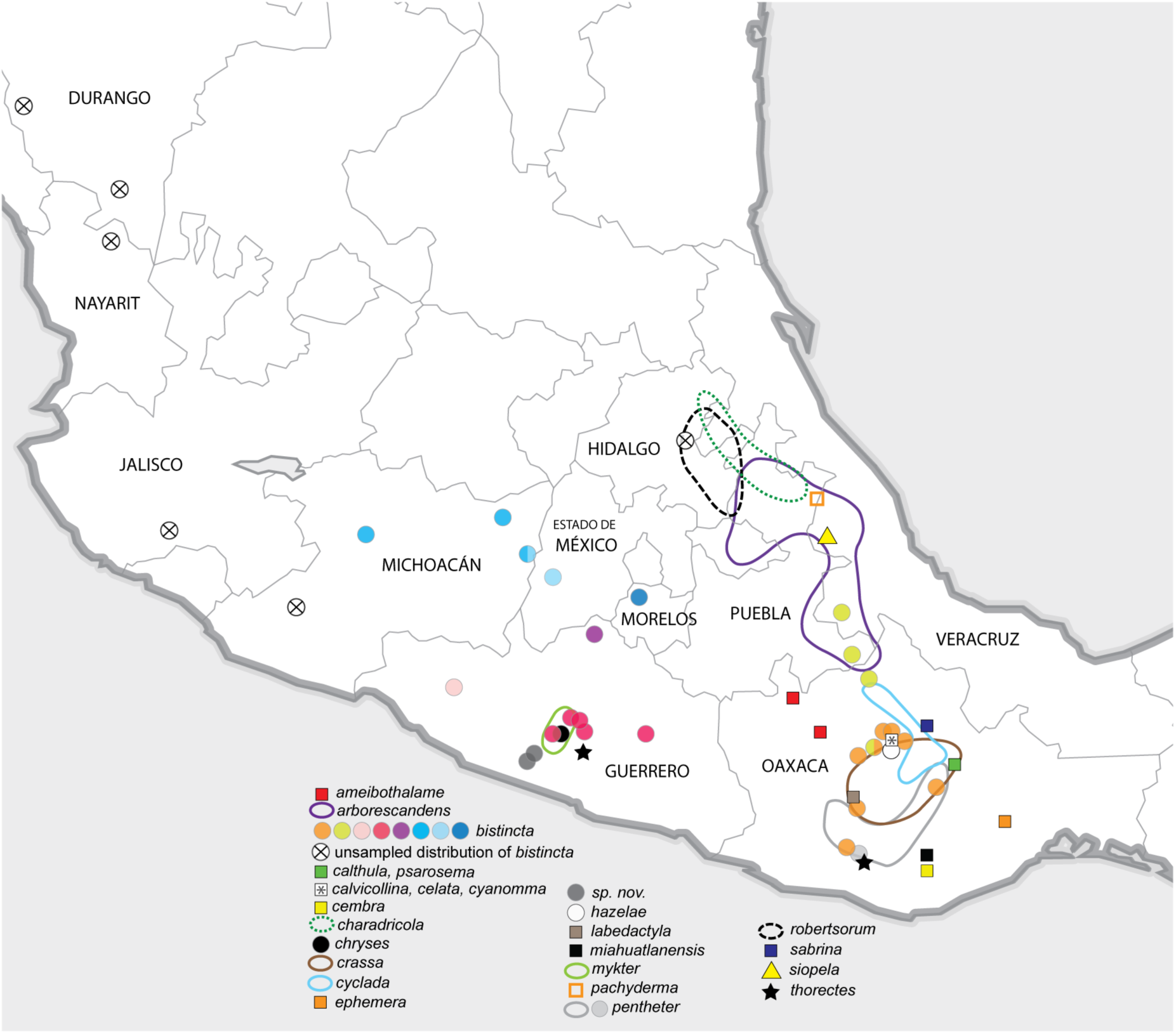
Sampled and unsampled parts of *S. bistincta* range in relation to known distributions (or localities, where distributional information is lacking) of other *Sarcohyla* species.

**Fig. S2.**
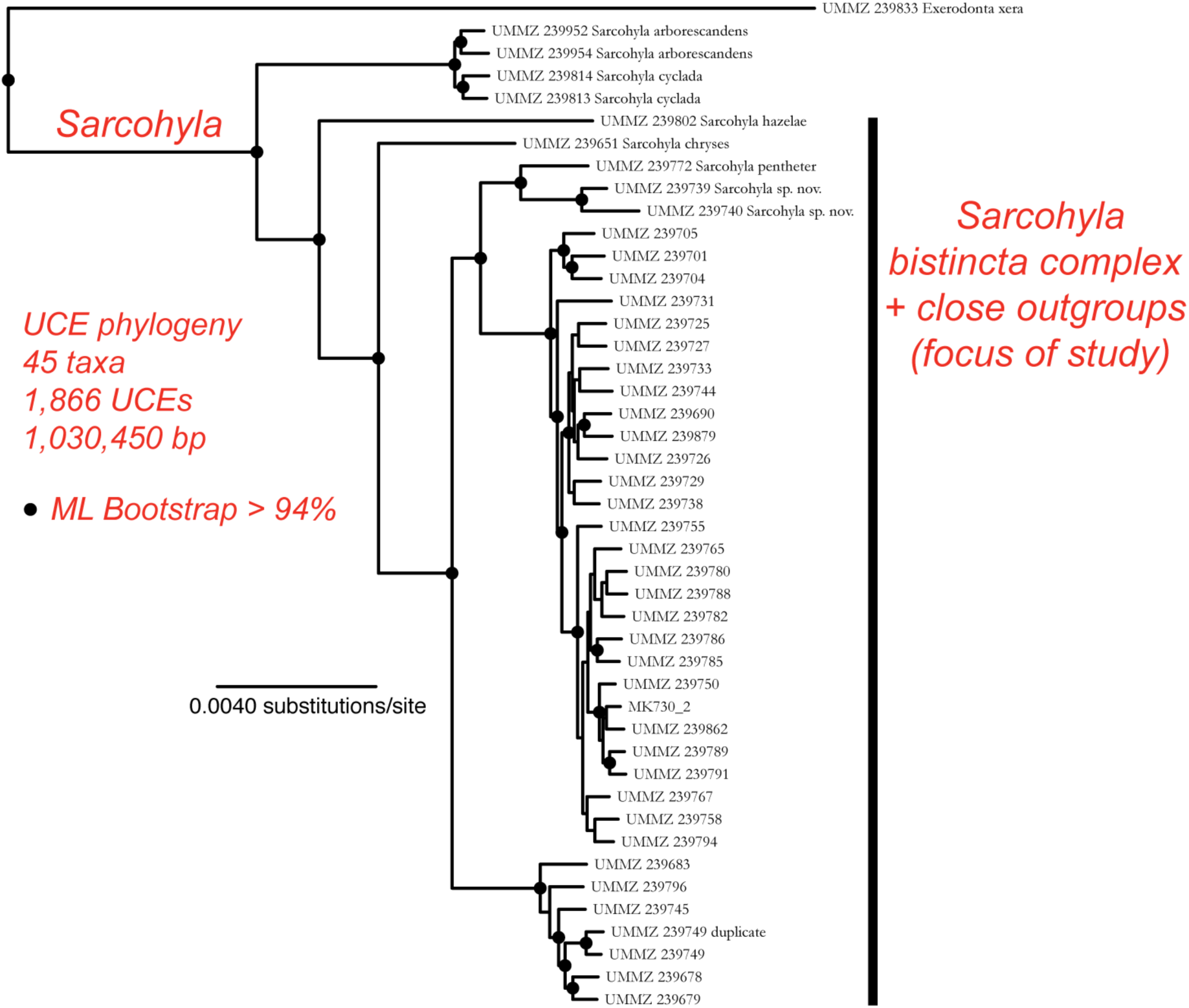
UCE tree of 45 samples of *Sarcohyla* and outgroup *Exerodonta xera* used to determine the ingroup.

